# Next-generation diagnostics: virus capture facilitates a sensitive viral diagnosis for epizootic and zoonotic pathogens including SARS-CoV-2

**DOI:** 10.1101/2020.06.30.181446

**Authors:** Claudia Wylezich, Sten Calvelage, Kore Schlottau, Ute Ziegler, Anne Pohlmann, Dirk Höper, Martin Beer

## Abstract

**Background:** The detection of pathogens in clinical and environmental samples using high-throughput sequencing (HTS) is often hampered by large amounts of background information, which is especially true for viruses with small genomes. Enormous sequencing depth can be necessary to compile sufficient information for identification of a certain pathogen. Generic HTS combining with in-solution capture enrichment can markedly increase the sensitivity for virus detection in complex diagnostic samples.

**Methods:** A virus panel based on the principle of biotinylated RNA-baits was developed for specific capture enrichment of epizootic and zoonotic viruses (VirBaits). The VirBaits set was supplemented by a SARS-CoV-2 predesigned bait set for testing recent SARS-CoV-2 positive samples. Libraries generated from complex samples were sequenced via generic HTS and afterwards enriched with the VirBaits set. For validation, an internal proficiency test for emerging epizootic and zoonotic viruses (African swine fever virus, Ebolavirus, Marburgvirus, Nipah henipavirus, Rift Valley fever virus) was conducted.

**Results:** The VirBaits set consists of 177,471 RNA-baits (80-mer) based on about 18,800 complete viral genomes targeting 35 epizootic and zoonotic viruses. In all tested samples, viruses with both DNA and RNA genomes were clearly enriched ranging from about 10-fold to 10,000-fold for viruses including distantly related viruses with at least 72% overall identity to viruses represented in the bait set. Viruses showing a lower overall identity (38% and 46%) to them were not enriched but could nonetheless be detected based on capturing conserved genome regions. The internal proficiency test supports the improved virus detection using the combination of HTS plus targeted enrichment but also point to the risk of carryover between samples.

**Conclusions:** The VirBaits approach showed a high diagnostic performance, also for distantly related viruses. The bait set is modular and expandable according to the favored diagnostics, health sector or research question. The risk of carryover needs to be taken into consideration. The application of the RNA-baits principle turned out to be user-friendly, and even non-experts (without sophisticated bioinformatics skills) can easily use the VirBait workflow. The rapid extension of the established VirBaits set adapted to actual outbreak events is possible without any problems as shown for SARS-CoV-2.

## Background

Disease control is encompassing several fields like powerful diagnostics for early detection, efficient therapy and disease prophylaxis as for example vaccination. The present study focused on broad and powerful diagnostics for viral pathogens including evolved, newly emerging or unrecognized viruses. The detection of the latter can be challenging for conventional virus diagnostics relying on specific quantitative PCR assays. Sequence information for such newly emerging pathogens is often rare or not existing. In addition, PCR-based diagnosis of co-infections with several pathogens is complicated, can be time-consuming and expensive, and therefore, co-infections can easily be overseen. Moreover, pathogen detection in cases of immunocompromised patients might be hampered due to a deviating course of the infection or its ectopic location [1]. For all the mentioned challenges, untargeted metagenomics using high-throughput sequencing (HTS) offers a swift and broad solution. It also enables the simultaneous detection of the genetic information of pathogens of all taxa including viruses, bacteria and eukaryotic pathogens like parasites or fungi [2]. However, poor sample quality, low pathogen loads and high levels of background consisting of nucleic acids of the host or accompanying bacteria often lead to a detrimental pathogen/background nucleic acid ratio of the final sequencing libraries [3]. Analyses of the resulting sequence datasets can be laborious and time consuming and difficult to interpret, even for experts. This is especially true for pathogenic viruses with rather small RNA genomes that can get lost in datasets generated from complex samples. Extremely large sequence datasets and intense time-consuming analyses would be necessary to compile enough meaningful information to finalize a genome of a certain pathogen [4]. This significantly impedes pathogen detection causing a diagnostic gap and making HTS difficult to implement in the daily routine of diagnostic laboratories.

The enrichment of certain pathogens prior to HTS using capture enrichment methods can help to increase the target sequence information considerably, along with avoiding extensive sequencing efforts, as recently reviewed by Gaudin and Desnues [3], and consequently, can enhance the sensitivity of diagnostic HTS. Basically, enrichment can be realized by the application of oligonucleotides (DNA probes or RNA baits) that are complementary to the sequence information of the target pathogens to reach a targeted positive separation of them. Target enrichment with complex samples was already applied for viral [5–7], bacterial [8], parasitical [9] and fungal [10] pathogens to compile full or nearly full genomes for follow-up analyses. For diagnostic reasons, however, the combination of probes or RNA baits of several pathogens into one set, depending on the scope, for example virus, microbiological or syndromic diagnostics, is beneficial. This approach can save numberless PCR reactions, which are specific for only one pathogen or pathogen groups. Studies implementing this practice mostly using DNA probes for different virus panels were already presented [11–13]. However, since DNA:DNA hybrids are less stable and have a weaker hybridization efficiency than RNA:DNA hybrids [14,15], RNA baits might be preferable.

For the purpose of improved virus diagnostics, we used RNA baits for an in-solution capture assay with subsequent HTS (**Fig. 1**). In a pilot study, we selected a panel of viral pathogens for notifiable epizootic and zoonotic diseases. A genome dataset for the relevant virus groups was created (containing about 18,800 full-length virus genomes) and a set of RNA baits (VirBaits) was derived from this dataset to facilitate the identification and subsequently diagnosis of viral pathogens in farm and reservoir animals but also humans. The resulting VirBaits set was tested for a set of representatives of pathogenic viruses with diagnostic samples from already examined cases instead of spiked samples. These clinical samples had, to some extent, challenging low virus titers. Due to the recent worldwide SARS-CoV-2 outbreak [16], we extended the bait panel for this newly emerging virus and successfully tested the set with SARS-CoV-2 positive samples.

**Figure 1.**
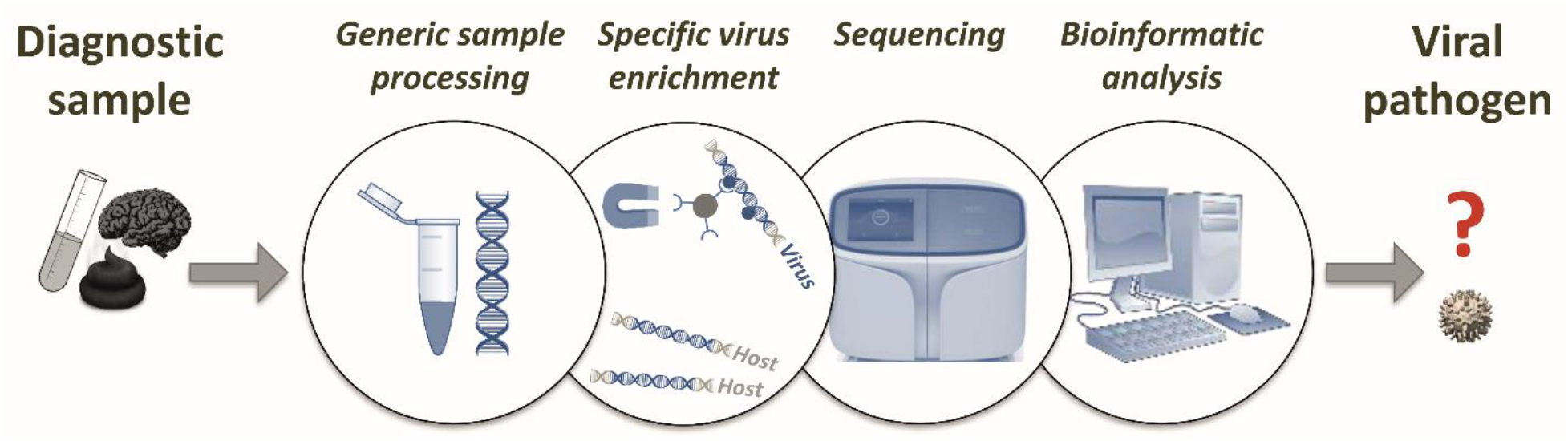
Sketch of the viral diagnostic HTS workflow combining generic metagenomics [2] with the VirBaits approach. Images used as symbols were obtained from the free website https://pixabay.com/.

## Methods

### Bait design for the targeted virus panel

The aim of the study was to design a diagnostic bait set for epizootic and zoonotic viral pathogens. For this purpose, virus genomes were compiled based on a curated list of complete virus genomes from the viralzone website (list “SEPT_2017” on viralzone website [17], comprising 70,352 virus genomes) for viruses causing notifiable epizootic and emerging zoonotic diseases. For Influenza A (IAV), a virus with RNA genome characterized by a broad genomic variety, all available IAV sequences covering full or nearly full-length segments from Influenza viruses collected from avian hosts that were available in FluDB June 2018 were taken (representing 12,608 isolates). Duplicated IAV sequences were removed and the resulting set of 79,859 segment sequences were used as basis for bait design. The final virus genome dataset comprising about 18,800 complete virus genomes for epizootic and zoonotic diseases (**Table 1**) was sent to Arbor Biosciences (Ann Arbor, Michigan, USA) for bait design. On this basis, 80-mer oligonucleotides were derived with 1.3× tiling density. They were afterwards checked against 19 possible host genomes (*Anas platyrhynchos, Anser cygnoides, Bos taurus, Camelus dromedaries, Canis familiaris, Capra hircus, Cervus elaphus, Chlorocebus sabaeus, Enhydra lutris, Equus caballus, Felis catus, Gallus gallus, Homo sapiens, Mus musculus, Ovis aries, Rattus norvegicus, Sus scrofa, Vicugna pacos, Vulpes vulpes*) to remove those oligonucleotides that were likely to hybridize and capture host nucleic acids. Finally, redundancy was decreased by removing oligonucleotides that were >95% identical to each other. Based on the final set, biotinylated RNA-baits were manufactured. The tailored custom myBaits kit (VirBaits) for 48 hybridization reactions including the RNA bait set comprising 177,471 baits and adapters blockers fitting with Ion Torrent sequencing adapters was obtained (Arbor Biosciences). All baits including the GenBank accession number of the respective virus genomes are provided in the supplementary file (fasta format).

**Table 1.**
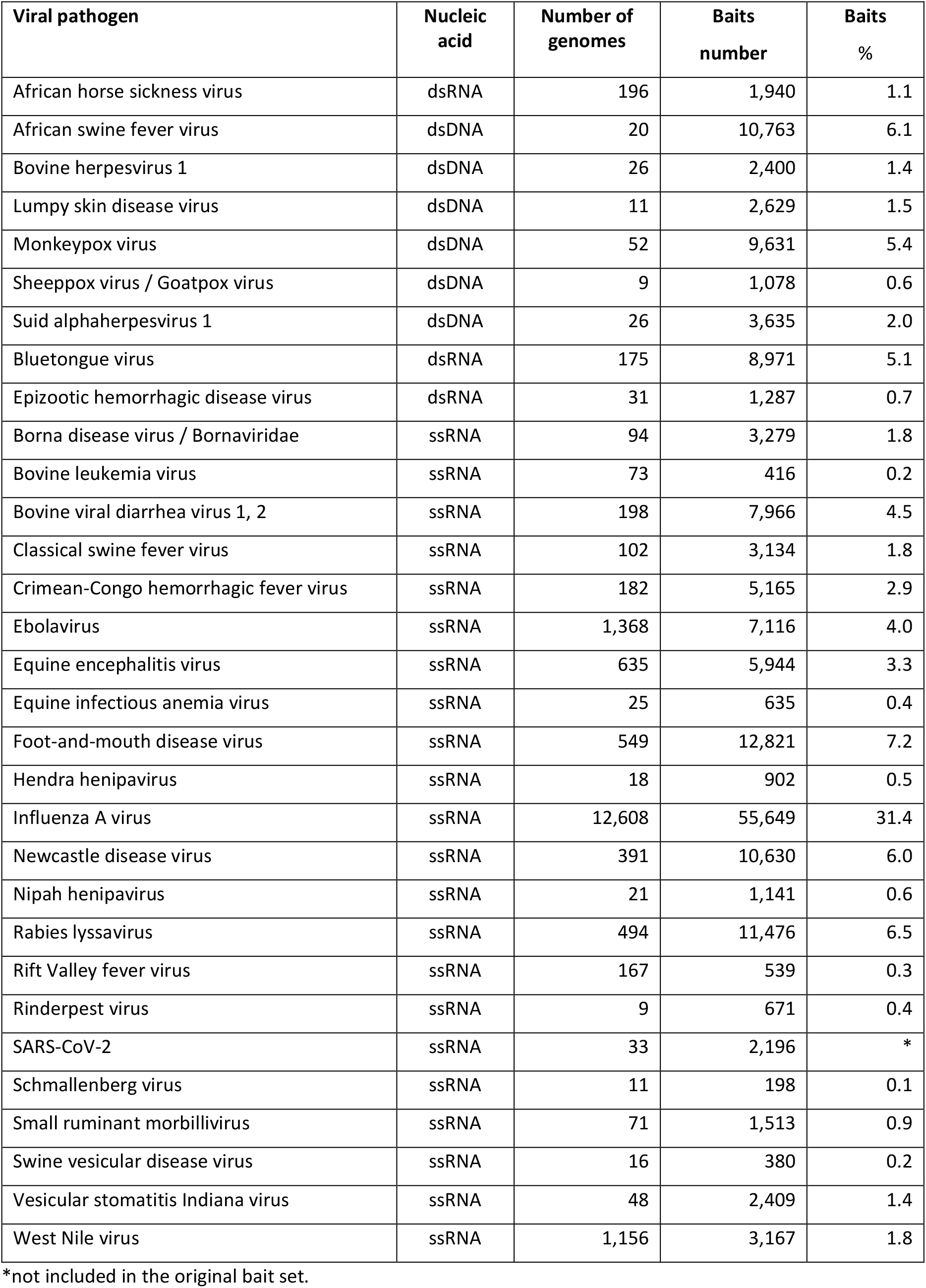
Viral pathogens causing notifiable or emerging epizootic and zoonotic diseases, their nucleic acids, number of genomes or genome segments and baits represented on the VirBaits set. The percentage of baits on the VirBaits set is given for each virus (compare **Figure 1**).

| Viral pathogen | Nucleic acid | Number of genomes | Baits number | Baits % |
| --- | --- | --- | --- | --- |
| African horse sickness virus | dsRNA | 196 | 1,940 | 1.1 |
| African swine fever virus | dsDNA | 20 | 10,763 | 6.1 |
| Bovine herpesvirus 1 | dsDNA | 26 | 2,400 | 1.4 |
| Lumpy skin disease virus | dsDNA | 11 | 2,629 | 1.5 |
| Monkeypox virus | dsDNA | 52 | 9,631 | 5.4 |
| Sheeppox virus / Goatpox virus | dsDNA | 9 | 1,078 | 0.6 |
| Suid alphaherpesvirus 1 | dsDNA | 26 | 3,635 | 2.0 |
| Bluetongue virus | dsRNA | 175 | 8,971 | 5.1 |
| Epizootic hemorrhagic disease virus | dsRNA | 31 | 1,287 | 0.7 |
| Borna disease virus / Bornaviridae | ssRNA | 94 | 3,279 | 1.8 |
| Bovine leukemia virus | ssRNA | 73 | 416 | 0.2 |
| Bovine viral diarrhea virus 1, 2 | ssRNA | 198 | 7,966 | 4.5 |
| Classical swine fever virus | ssRNA | 102 | 3,134 | 1.8 |
| Crimean-Congo hemorrhagic fever virus | ssRNA | 182 | 5,165 | 2.9 |
| Ebolavirus | ssRNA | 1,368 | 7,116 | 4.0 |
| Equine encephalitis virus | ssRNA | 635 | 5,944 | 3.3 |
| Equine infectious anemia virus | ssRNA | 25 | 635 | 0.4 |
| Foot-and-mouth disease virus | ssRNA | 549 | 12,821 | 7.2 |
| Hendra henipavirus | ssRNA | 18 | 902 | 0.5 |
| Influenza A virus | ssRNA | 12,608 | 55,649 | 31.4 |
| Newcastle disease virus | ssRNA | 391 | 10,630 | 6.0 |
| Nipah henipavirus | ssRNA | 21 | 1,141 | 0.6 |
| Rabies lyssavirus | ssRNA | 494 | 11,476 | 6.5 |
| Rift Valley fever virus | ssRNA | 167 | 539 | 0.3 |
| Rinderpest virus | ssRNA | 9 | 671 | 0.4 |
| SARS-CoV-2 | ssRNA | 33 | 2,196 | * |
| Schmallenberg virus | ssRNA | 11 | 198 | 0.1 |
| Small ruminant morbillivirus | ssRNA | 71 | 1,513 | 0.9 |
| Swine vesicular disease virus | ssRNA | 16 | 380 | 0.2 |
| Vesicular stomatitis Indiana virus | ssRNA | 48 | 2,409 | 1.4 |
| West Nile virus | ssRNA | 1,156 | 3,167 | 1.8 |

\*not included in the original bait set.

### Hybridization and enrichment procedure

The VirBaits set was applied according to the manufacturer’s instructions (myBaits manual v.4.01, Arbor Bioscineces, April 2018). In brief, hybridization reagents including baits were thawed, collected and mixed with each other preparing HYB and blockers master mixes. The HYB mix was incubated for 10 min at 60°C to clear the solution. After adapting to room temperature for 5 min, the HYB mix was aliquoted to 18.5 μl per reaction. The blockers mix was made and aliquoted to 5 μl each and subsequently combined with up to 7 μl of the sequencing library (resulting in the LIB mix). The LIB mix of each library was denatured for 5 min at 95°C using a thermal cycler. Both, the LIB and HYB mixes were afterwards incubated for 5 min at 65°C. Subsequently, 18 μl of the HYB aliquot were mixed with the LIB mix aliquot each (12 μl) using pipetting up and down and spinning down, and then incubated for about 16 h at the suggested hybridization temperature (65°C). We tested three variations from the protocol recommended in the manual. (i) The Rabies virus (RABV) infected sample L03007 was hybridized with only the half amount of the reaction volume. (ii) For IAV-infected samples L02651 and L03706, the baits in the HYB mix were diluted 1:5 with ultrapure water. (iii) The Kotalathi Bat lyssavirus infected sample L02374 was incubated at 62°C instead at the default hybridization temperature of 65°C. After hybridization, the samples were purified three times at the hybridization temperature using binding beads contained in the kit and finally eluted with 30 μl 10 mM Tris-Cl, 0.05% Tween-20 solution (pH 8.0-8.5). An aliquot (16 μl) of the enriched samples was amplified using the GeneRead DNA Library L amplification Kit (Qiagen) according to manufacturer’s instructions performing 14 cycles and purified using solid-phase paramagnetic bead technology. Quality check and quantification of the enriched libraries was performed as described [2].

### Direct comparison of bait-treated with untreated sequencing datasets

The performance of the VirBaits set was tested for selected viruses including such with RNA and DNA genomes. We mainly used infected sample material of real cases that had already been sequenced for diagnostic purpose but also a spiked salmon sample from a laboratory proficiency test (**Table 2**). The libraries of those samples were generated using a validated unbiased workflow [2] without any enrichment or depletion. In the present study, these previously constructed and archived sequencing libraries (stored at −20°C) were treated with the bait set as described above. Sequence datasets obtained with and without VirBaits treatment were analyzed via reference mapping using a reference genome of the respective virus and the Genome Sequencer software suite (versions 2.6; Roche) to directly compare the containing virus amount in terms of virus reads of the datasets. The enrichment factor was calculated by dividing the percentage of enriched virus amount by original virus amount for each sample. Most tested viruses were nearly identical or closely related to the viruses in the underlying genome set used for the bait design (case 1 samples). Beside these viruses, we also tested samples infected with viruses that indeed belong to the virus families included in the bait design but that are only distantly related to the sequences used for the bait design (**Table 3**). The taxonomic composition of the VirBaits metagenomics datasets were additionally analysed using the RIEMS tool [18].

**Table 2.** Overview of tested libraries: case 1 samples. For viruses enclosed in the samples, reference genomes were included in the genome set used for VirBait design. Sample matrix, animal species, the detected virus and the references are given. The amount of virus reads in the sequence datasets found in the originally sequenced libraries and in the VirBaits treated libraries is given as percentage. Libraries were made from RNA except the one marked by an asterisk, which was made from DNA. # data set size is given in terms of reads.

| Library ID | Sample material | Verified virus | Virus portion |  | Data set size<br>*VirBaits*<sup>#</sup> | Reference |
| --- | --- | --- | --- | --- | --- | --- |
|  |  |  | *generic HTS* | *VirBaits* |  |  |
| L02725 | Brain, *Strix nebulosa* | WNV | 0.14% | 58.7% | 2,678 | [28] |
| L02726 | Spleen, *Strix nebulosa* | WNV | 5.6% | 91.4% | 3,857 | [28] |
| L03224 | Spinal cord, *Equus* sp. | WNV | Not sequenced | 0.2% | 1,305,078 | unpublished |
| L03038 | Brain, *Bubo scandiacus* | WNV | 0.0003% | 0.03% | 155,648 | Santos et al., in preparation |
| L03451 | Spinal cord, *Equus* sp. | WNV | 0.0006% | 0.2% | 907,780 | [23] |
| L02838 | Brain, *Canis lupus* | RABV | 0.002% | 7.9% | 158,514 | unpublished |
| L03007 | Brain, *Tragelaphus* sp. | RABV | 1.5% | 97.5% | 890,113 | unpublished |
| L03015 | Salivary gland, *Tragelaphus* sp. | RABV | 0.005% | 45.8% | 1,060,955 | unpublished |
| L02912 | Brain, *Rattus* sp. | BoDV | 0.75% | 85.2% | 183,579 | Schlottau et al., in preparation |
| L02913 | Brain, *Rattus* sp. | VSBV-1 | 0.0004% | 0.6% | 42,793 | Schlottau et al., in preparation |
| L02456 | Soft Palate Cells, *Bos taurus* | FMDV | 0.002% | 2.0% | 719,153 | [30] |
| L02568* | Food matrix spiked with a mock community | BHV | 0.0003% | 0.04% | 181,338 | [32] |
| L02569 |  | BVDV | 0% | 0.007% | 506,465 |  |
|  |  | BDV | 0.00001% | 0.01% |  |  |
| L02762* | Spleen, *Sus scrofa* | ASFV | 0.05% | 20.7% | 1,353,258 | unpublished |
| L02651 | Organ pool, *Gallus gallus* | IAV (H5N8) | 0.01% | 70.1% | 280,247 | unpublished |
| L03706 | Brain, *Gallus gallus* | IAV (H3N1) | 0.5% | 77.0% | 317,862 | unpublished |
| L03827 | Nasal wash, *Mustela putorius* | SARS-CoV-2 | 0.3% | 53.8% | 1,248 | [31] |
| L03828 | Nasal wash, *Mustela putorius* | SARS-CoV-2 | 0% | 9.1% | 449 | [31] |
| L03829 | Nasal conchae, *Mustela putorius* | SARS-CoV-2 | 0.00004% | 0.001% | 194,891 | [31] |
| L03830 | Nasal conchae,<br>*Mustela putorius* | SARS-CoV-2 | 0.02% | 14.9% | 325,694 | [31] |
| L03862 | Nasal conchae,<br>*Mustela putorius* | SARS-CoV-2 | 0.0004% | 0.2% | 960,594 | [31] |
| L03862 | Nasal conchae,<br>*Mustela putorius* | SARS-CoV-2 | 0.001% | 0.5% | 946,268 | [31] |
| L03862 | Brain, *Mustela putorius* | SARS-CoV-2 | 0,00002% | 0,005% | 294,146 | [31] |

**Table 3.** Overview of tested libraries: case 2 samples. For the viruses enclosed in the samples, for that no reference genomes were included in the genome set used for the VirBaits design. Sample matrix, animal species, the detected virus and the references are given. The amount of virus reads in the sequence datasets found in the originally sequenced libraries and in the VirBaits treated libraries is given as percentage. Libraries were made from RNA except the one marked by an asterisk, which was made from DNA.

| Library ID | Sample material | Verified virus | Generic HTS |  | VirBaits |  | Reference |
| --- | --- | --- | --- | --- | --- | --- | --- |
|  |  |  | Total reads | Virus portion | Total reads | Virus portion | Reference |
| L03038 | Brain, *Bubo scandiacus* | USUV | 4,347,713 | 0.001% | 155,648 | 0.1% | Santos et al., in preparation |
| L02262 | Spinal cord, *Ovis aries* | OvPIV | 2,422,322 | 0.06% | 261,673 | 0.04% | [19] |
| L01003* | Cell culture supernatant | PeHV | 982,587 | 0.6% | 154,820 | 0.4% | [21] |
| L02374 | Organ suspension, bat | KBLV | 5,696,571 | 3.4% | 508,034 | 89.6% | Freuling et al., in preparation |

### Variant analysis

Variant analyses were performed to compare within-sample variant frequencies in the RABV genome of sample L03007 and the KBLV genome of sample L02374 before and after VirBaits treatment. The whole virus genomes of RABV and KBLV were received by mapping the sequencing datasets against the lyssavirus reference sequences (NC_001542.1), and by de-novo assembly and subsequent mapping, respectively, using the Genome Sequencer software suite (version 3.0; Roche). Subsequently, the obtained whole genome sequence was used in a second mapping round as reference sequence in order to calculate the total number of viral reads for the respective datasets. To identify potential single nucleotide variants (SNPs), the Torrent Suite plugin Torrent variantCaller (version 5.12) was used (parameter settings: generic, S5/S5XL(530/540), somatic, low stringency, changed alignment arguments for the TMAP module from map 4 [default] to map1 map2) while the previously determined whole genome sequence was set as reference. In addition, the variant analysis tool integrated in Geneious Prime (2019.2.3) was applied for further confirmation and the acquisition of variant frequencies for each variant (default settings, minimum variant frequency 0.02). Positions of identified potential SNPs were visually inspected in Geneious Prime.

### Proficiency testing of the VirBaits application

The broad functionality of the VirBaits approach was investigated via an internal proficiency test using five blind samples. The test initiator (M.B.) provided five samples (see **Table 4**) only declaring sample number and type (tissue sample with Trizol or already extracted RNA) whereas the sample processor (C.W.) did not get any information on the samples like host, virus, kind of organ or tissue or pre-diagnosis.

**Table 4.** Proficiency test samples with information on viruses contained and found after VirBaits treatment and dataset size.

| Sample number | Sample matrix provided | Sample details | Total reads | Virus contained | Virus found *VirBaits* | Virus portion |
| --- | --- | --- | --- | --- | --- | --- |
| X1 | Animal sample with Trizol | Spleen of ASFV-infected wild boar | 644,441 | ASFV | ASFV | 0.01% |
| X2 | Animal sample with Trizol | Saliva sample of FMDV-infected cattle | 23,870 | FMDV | FMDV | 72% |
| X3 | Extracted RNA | RVFV vaccine strain MP-12 with Vero 76 cells | 24,022 | RVFV | RVFV | 98% |
|  |  |  |  |  | NiV | 0.5% |
| X4 | Extracted RNA | Mix of RNA (10 µl each) of Ebolavirus, Marburgvirus, Nipah henipavirus | 256,408 | NiV | NiV | 45% |
|  |  |  |  | EOV | EOV | 0.6% |
|  |  |  |  | Marburgvirus | Marburgvirus | 0.05% |
| X5 | Cells with Trizol | African green monkey kidney cells | 269,701 | None | None | 0 |

### Extension of the VirBaits set: Tests with SARS-CoV-2 positive samples

To expand the above-mentioned VirBaits set for the emerging SARS-CoV-2 virus, a predesigned bait set for the coronavirus SARS-CoV-2 was designed and available free of charge from Arbor Biosciences. This SARS-CoV-2 bait set was mixed with the VirBaits set in the ratio of 1:40 to simulate an extended VirBaits set, and applied as described above. Different samples of SARS-CoV-2 infected ferrets (**Table 2**; Schlottau et al. submitted) were used for tests with the extended VirBaits set.

## Results

### Design of the virus in-solution capture assay (VirBaits)

The VirBaits panel covered 35 viral pathogens of notifiable epizootic and zoonotic diseases (**Table 1**). The resulting bait set comprises 177,471 biotinylated RNA-baits complementary to 18,800 virus genomes (**Fig. 2**). The final number of baits in the VirBaits set among the target virus groups depends on the diversity of viruses comprised in the underlying genome set, the number of genomes and the corresponding genome size, respectively. The highest portion of baits belongs to Influenza A viruses that have small segmented single-strand RNA genomes with a broad variety of genetic information that needs to be covered. Therefore, 79,859 Influenza A segments sequences from 12,608 Influenza isolates were used for bait design and resulted in 55,649 baits that comprise 31.4% of the entire set. In contrast, the smallest portion of only 11 underlying complete genomes for Schmallenberg virus (SBV) with a small, segmented single-strand RNA genome with limited genomic variety resulted in 198 baits (0.1% of the entire set).

**Figure 2.**
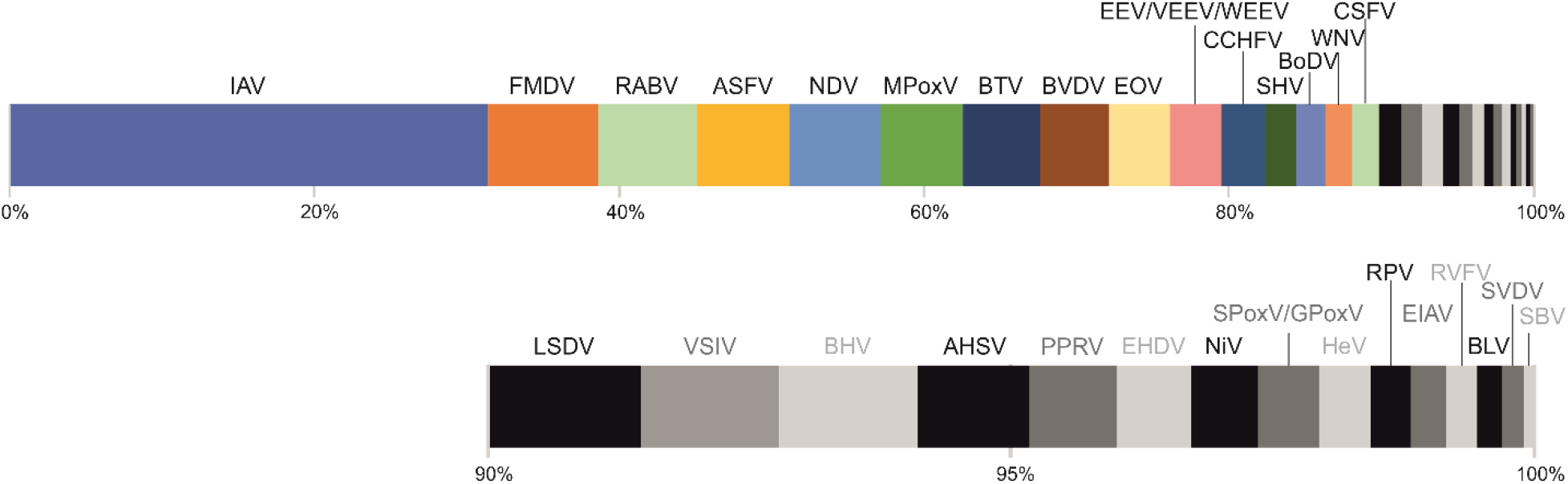
The distribution of RNA-baits among the virus groups included in the final VirBaits panel is shown. The black-grey part of the upper panel is zoomed out in the lower panel.

### Testing of the VirBaits set by directly comparing generically generated and bait-treated sequencing datasets

The VirBaits performance was assessed via direct comparison of datasets generated with an untargeted HTS approach for diagnostic reasons and datasets additionally generated from the same archived libraries after treatment with the VirBaits set. The portion of virus reads extracted from the metagenomics datasets using reference mapping for both datasets per sample and the enrichment factor are shown (**Table 2**, **Figure 3**). In general, the effect of the VirBaits treatment was very high. Two cases can be distinguished: viruses, which are comprised in the bait set and viruses of which only ditant relatives are comprised in the bait set.

**Figure 3.**
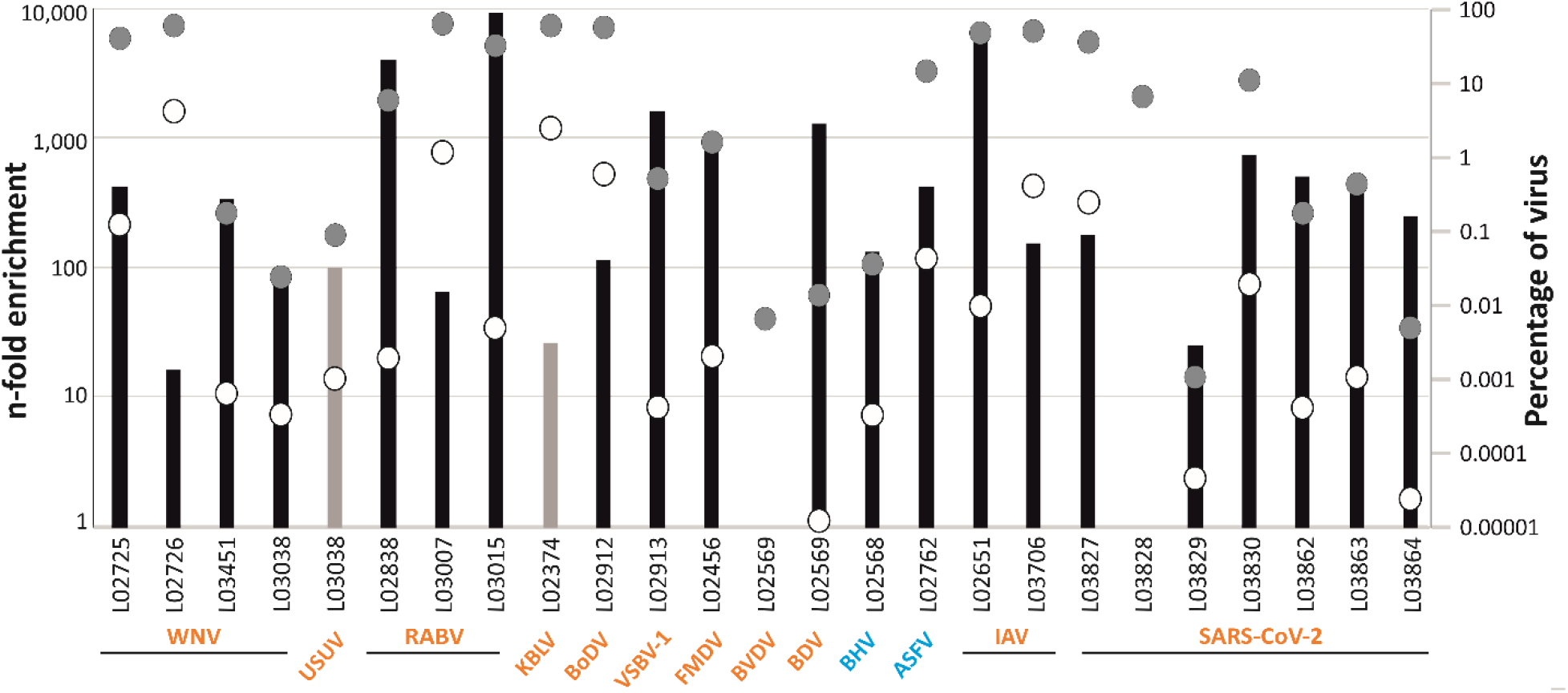
Enrichment of virus reads after VirBaits treatment. The numbers refer to the tested samples according to **Table 2** and **Table 3**. Black and grey bars indicate viruses that were included or not included in the bait design, respectively (labelling at left axis). White and grey dots indicate the virus amount (percentage) in datasets obtained without and with VirBaits treatment, respectively (labelling at right axis). RNA viruses are marked in orange; DNA viruses are marked in blue.

Case one, for all tested viruses that are represented in the VirBaits set, we could reach a clear enrichment from 16-fold for sample L02726, slightly loaded with West Nile virus, to 9,200-fold for the Rabies lyssavirus infected sample L03015 (**Table 2, Figure 3**). Also for viruses with large DNA genomes (i.e. African swine fever virus sample L02762, Bovine herpesvirus 1 sample L02568), a clear increase of virus reads could be detected after the VirBaits treatment. Likewise, viruses that are only less represented in the bait set (e.g., West Nile virus 1.8%, Bovine herpesvirus 1.4%, **Table 1**) were properly enriched (**Figure 3**). The additionally performed tests with the complemented SARS-CoV-2 set resulted in a clear increase of coronavirus reads (9 to 745-fold). Nearly complete virus genomes can be assembled from many samples including samples with a high host genome background and low-level viral genome loads.

Case two, the results vary regarding samples infected with viruses distantly related to the viruses included in the underlying genome set, although related taxa of the same virus family are represented in the VirBaits set. On the one hand, we found a clear enrichment for two samples: sample L02374 infected with KBLV (Kotalathi bat lyssavirus, showing an overall identity of 72% to RABV genomes) was enriched 26-fold, and sample L03038 infected with Usutu virus (showing about 74% overall identity to WNV; compare **Table 3** and **Fig. 3**) was enriched 100-fold. On the other hand, two samples did not show any increase after VirBaits treatment but were at least detectable. These are sample L02262 infected with a novel ovine picornavirus (showing about 46% and 38% overall identity to SVDV and FMDV genomes in the VirBaits set, respectively) and sample L01003 infected with a penguin alphaherpesvirus (showing about 38% and 36% overall identity to SHV and BHV genomes, respectively; compare **Table 3**). The genome coverage for those samples shows a distribution of reads over the complete genome (L02374, L03038) or the concentration in certain conserved regions (L01003, L02262). Beside the targeted viral reads, the datasets mainly contained host-related reads (Mammalia) and some bacterial reads including contaminations as typical for the used sample processing workflow [2].

### Analysis of variants in lyssavirus genomes

For sample L03007, the complete Rabies lyssavirus genome assembly generated via generic HTS did show the same single nucleotide variants (SNV) as the genome assembly of the virBaits dataset. The variants in each genome assembly were found to exhibit nearly unaltered frequencies per SNV (**Fig. 4**) indicating an equal extraction of Rabies virus reads using the bait treatment. The same is true for the KBLV-infected sample L02374 showing two point mutations with similar variant frequencies.

**Figure 4.**
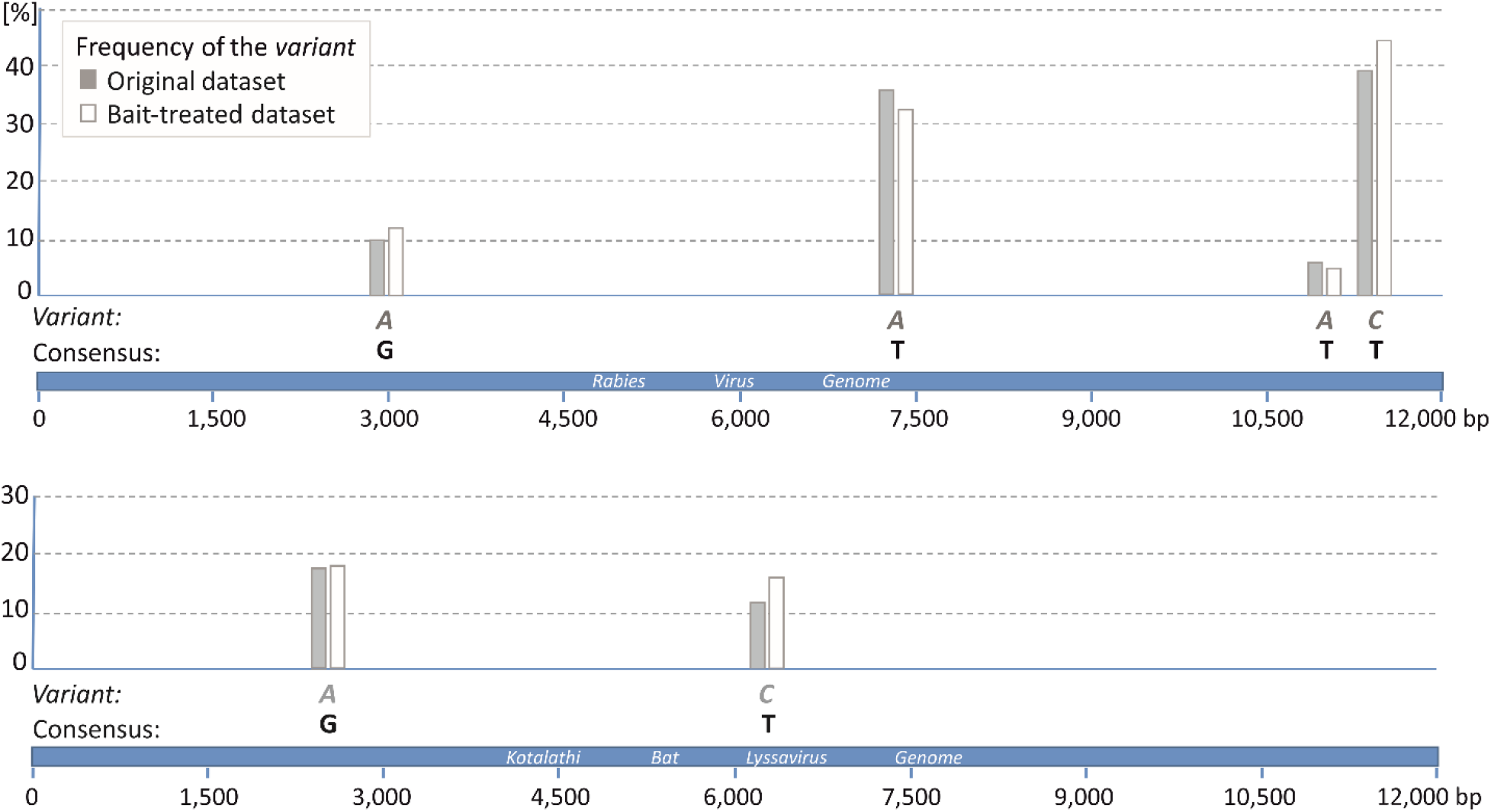
Schematic representation of variant analysis results for the brain sample L03007 infected with RABV (upper panel) and sample L02374 infected with KBLV (lower panel). Portions of the variants are given for the dataset processed with the untargeted workflow (grey bars) and for the dataset after VirBaits treatment (white bars). Four and two point mutations deviating from the consensus were detected along the RABV genome (at positions 2,926, 7,419, 11,245 and 11,790) and the KBLV (at positions 2,547 and 6,172), respectively, by applying a strand bias less than 70%.

### Testing of the VirBaits set with blind samples – proficiency test

In the proficiency test blind samples, high impact transboundary or zoonotic viruses detected in the samples X1 to X4 were ASFV, FMDV, RVFV, NiV, EOV and Marburgvirus (**Table 4**). In sample X5, no virus but instead reads related to the green monkey cell line was detected. The assignments made by the sample processor (C.W.) were approved as correct by the proficiency test initiator (M.B.). However, in sample X3, a small amount of NiV was detected that was not supposed to be in there indicating a carryover from sample X4 indeed containing NiV. Using mapping analyses of all samples against all viruses that were included in the proficiency test no other indication of carryover was found in any other proficiency test dataset.

## Discussion

The aim of the present study was the design and test of a virus enrichment panel, whose application markedly enhances the virus signal in diagnostic metagenomics by targeted capture enrichment. In general, the VirBaits approach with custom-designed kits using 80-mer RNA probes was easy to apply for our purpose by just compiling the relevant virus genomes and providing them to the supplier. In our opinion, it is applicable for diagnosticians who are in need for bait sets to capture specific pathogens but are not proficient in bioinformatics skills implementing the underlying genomes into meaningful oligonucleotides.

As a base for this HTS plus capture enrichment approach, we used a generic HTS workflow without applying rRNA depletion or PCR-amplification. This procedure ensures that the resulting sequencing data reflect the original sample composition in the best way [2]. This generic HTS approach is a powerful tool and led to the detection of novel viruses in many cases [19–22], although the initial presence of the virus can be low in some cases [19,22,23]. Indeed, samples with very low pathogen (virus) content represent a typical diagnostic setting. Therefore, samples with a range of genome loads including very low virus titers were used in the present pilot study to mimic the realistic clinical situation.

The designed VirBaits panel was successfully applied for all tested viruses, especially for viruses that are represented in the underlying genome set (case 1 samples). Altogether, an increase of 9-fold to 9,200-fold of the target sequence reads was achieved. When looking at the less enriched samples (**Fig. 3**), it is apparent that most of them show a very high percentage of virus amount in the VirBaits dataset indicating a certain saturation of the hybridization. Hence, a much higher increase would not be possible for these samples. Consequently, especially the low loaded samples benefit from the capture enrichment procedure. According to the myBaits manual, it is highly recommended to amplify libraries before enrichment. However, since we always use a PCR-free generic HTS workflow, we did not follow this recommendation and still obtained good results. This is in agreement with Mamanova et al. [24], who determined a severe bias when doing a PCR step before hybridization capture. The authors recommend keeping library amplification to a minimum and only performing 14-18 cycles after the capture hybridization especially with clinical samples exhibiting nucleic acids of lower integrity. The virome capture sequencing approach for vertebrate viruses, VirCapSeq-VERT [12], relies on sample pre-treatments for virus enrichment like filtration of samples, treatment of samples with ribonucleases to reduce extra-viral RNA and subsequent depletion of ribosomal RNA. Our study clearly shows that an amplification- and depletion-free HTS workflow is well suitable to be combined with capture enrichment. In addition, the application of RNA-baits instead of DNA probes leads to a high increase of the target viruses including moderately related viruses (case 2 samples). However, care must be taken when working with RNA baits, which are not as stable as DNA probes, by avoiding contaminations with RNases during handling and assuring the cooling chain. The bait amount per virus group in the entire panel was varying, yet the performance for virus groups that are represented by few baits was satisfying as shown for RVFV, BHV, HeV and NiV (**Fig. 3**, **Table 4**) indicating a robust method with high efficiency. Regarding samples infected with the OvPiV and the PeHV, the viruses are only distantly related to the viruses in the underlying genome set (case 2 samples). For such cases, an adaption of hybridization temperature and time might increase the output for those distantly related viruses. For the latter modification, evaporation might become a serious problem that needs to be considered. Nonetheless, distantly related viruses are at least detectable by the standard VirBaits procedure via capturing conserved regions. Hence, they would not get lost during the procedure. Altogether, the results indicate a high profitableness for all samples including low-load virus samples if included in the VirBaits set or moderately related to those viruses.

The tiling density is low in the present VirBaits panel (1.3×) compared with panels for single species applying a tiling density of 3× [6] or 4× [25]. This is owing to the diagnostic approach we were intending to design, improving diagnostic sensitivity but not necessarily leading to the assembly of whole genomes. Typically, one specific read in a sequence dataset indicating a pathogenic virus would be enough to call a suspicion and trigger follow-up analyses. However, as shown here, the used tiling density in many cases allows for the assembly of nearly complete genomes emphasizing the sensitivity of the approach. In addition, the detected SNVs of RABV and KBLV genomes point to equal frequencies (**Fig. 4**) indicating a consistent recovery of the genomes via capture enrichment as already demonstrated by Metsky et al. [13].

A general benefit of capture approaches is that no prior decision is necessary for which pathogen (virus) is sought after, i.e., which specific test system needs to be applied. This saves time and money by avoiding application of many laborious diagnostic techniques successively until a causative agent can be suspected and confirmed. Moreover, a negative result of a PCR approach can mask an undetected viral pathogen when PCR primers do not fit anymore with the genome of evolved pathogens. Otherwise, with positive PCR result, co-infections with several different pathogens can be overseen. For cases with atypical clinical signs or deviating viral pathogens, it might be that the relevant specific diagnostics are not considered or not existing, respectively. The synchronous detection of different viruses, as shown for sample L03038 having a co-infection with WNV and USUV, enabled by the generic HTS approach in combination with VirBaits approach is a great benefit. Compared to multiplex PCR approaches, that allow the detection of several viruses or virus variants in parallel [26,27], the VirBaits approach has a broader applicability and often provides sequence information of full or nearly full genomes instead of small gene fragments. This information can additionally be used for follow-up analyses on genetic diversity, virulence or functional potential.

The results of the internal proficiency test corroborate the applicability of the virBaits workflow combined with HTS as all-in-one approach for different pathogenic viruses without the necessity to do laborious diagnostic tests for at least each single virus or small groups of viruses. The datasets generated were relatively small but resulted in all cases in enough to determine the respective pathogen without any doubt. This also indicates the potential for reduction of sequencing costs when applying an enrichment method. For sample X1 infected with the DNA virus ASFV, the portion of virus reads in the dataset was only low since the sequencing library was made from RNA according to the standard HTS workflow [2]. In that case, starting from DNA would have delivered much higher read numbers, as seen for sample L02762 yielding a proper amount of African swine fever virus reads (compare **Table 2**). In sample X3, some reads of the NiV were found while they were not supposed to be in this sample but only in sample X4. This point to the risk of carryover that can be an issue for manual sample handling. Carryover contaminations typically occur in laboratories but become especially visible when highly enriched samples are used, like amplicons, or bait-enriched samples. An automated procedure or a cartridge-based system may avoid carryover in the future by excluding the “human factor”.

The rapid inclusion of baits and samples related to the recent SARS-CoV-2 outbreak demonstrate the high dynamics and ad-hoc adaptability of the here tested approach. The results for those samples show that the VirBaits workflow can be updated very rapidly and easily for viruses causing newly emerging health threats. In addition, we investigated several samples related to the WNV epidemic in Germany [23,28] including two horse samples. Horses are dead-end hosts for WNV and horse samples show generally lower virus titers in contrast to samples of bird being reservoir hosts for WNV infection. As shown here, the VirBaits treatments considerably improved the WNV detection for horse samples (L03451).

The application of myBaits is still expensive, although sequencing costs decrease with this procedure since no large data sets are necessary (compare **Tables 2-4**). The reduction of the reaction volume by 50%, as shown here for L03007, can reduce the cost for the enrichment reagents. Nevertheless, this opportunity needs to be validated in further trials and might depend on the individual sample type and the contained virus.

Resulting from the present pilot study, the VirBaits panel can be improved by optimizing the distribution of baits per virus group. This especially means a significant reduction of IAV baits, as shown here with the dilution of the baits for IAV samples, but also other viruses with small RNA genomes like FMDV, RABV, NDV (compare **Fig. 2**). Redundancy of the baits can be decreased by removing oligonucleotides that were >90% identical to each other (in contrast to 95% as used for the pilot study). This decrease should still allow good hybridization results considering the performance of deviating baits as shown here for KBLV, USUV, OvPiV and PeHV. Pooling of multiple libraries (for cost reduction), application of reduced hybridization temperatures or prolonged hybridization times (up to 48 hours) need to be investigated for distantly related targets. The latter step has to go along with procedures to avoid strong evaporation holding the reaction volume nearly constant during extended incubation time.

## Conclusions

The biotinylated RNA-baits principle for capture enrichment with a large and diverse virus panel as tested here is highly suitable to improve diagnostics for viruses causing notifiable epizootic and zoonotic diseases. It can be dynamically updated with new RNA-baits because of the modular concept that allows the addition of new bait sets for emerging diseases. The combination of a generic and manipulation-free HTS workflow with the capture approach leads to an enhancement of the viral signal in terms of sequence reads in the sequence dataset resulting in higher sensitivity of HTS. The identification of distantly related viruses can be difficult as shown for a deviating avian bornavirus in a proficiency test for HTS bioinformatics [29]. Although the problem in the mentioned study was related to bioinformatics analyses, the accumulation of certain virus reads by capture enrichment can avoid overlooking pathogenic viruses contained in a sample. The application of such an approach is recommendable for cases of unclear clinical pictures and can replace time-consuming screenings using large panel PCR diagnostics for specific single pathogens. This leads to an accelerate diagnosis and a very high detection rate. Although not the primary intention, the application of the VirBaits even enables the generation of nearly complete genome sequences in most cases.

## Supplementary information

Supplementary information accompanies this paper at {*link to be included upon acceptance*}.

### Additional file 1

Baits; all baits including the GenBank accession number of the respective virus genomes are provided (fasta format, 26 kb).

## Abbreviations

HTS: high-throughput sequencing
rRNA: ribosomal RNA
dsDNA: doubles-strand DNA
dsRNA: doubles-strand RNA
ssRNA: single-strand RNA

## Viruses

AHSV: African horse sickness virus
ASFV: African swine fever virus
BLV: Bovine leukemia virus
BHV: Bovine herpesvirus 1
BoDV: Borna disease virus
BTV: Bluetongue virus
BVDV: Bovine viral diarrhea virus 1, 2
CCHFV: Crimean-Congo hemorrhagic fever virus
CSFV: Classical swine fever virus
EEEV/VEEV/WEEV: Eastern, Venezuelan, and Western Equine encephalitis virus
EHDV: Epizootic hemorrhagic disease virus
EIAV: Equine infectious anemia virus
EOV: Ebolavirus
FMDV: Foot-and-mouth disease virus
HeV: Hendra henipavirus
IAV: Influenza A virus
KBLV: Kotalathi bat Lyssavirus
LSDV: Lumpy skin disease virus
MPoxV: Monkeypox virus
NDV: Newcastle disease virus
NiV: Nipah henipavirus
OvPiV: Ovine picornavirus
PeHV: Penguin alphaherpesvirus
PPRV: Small ruminant morbillivirus
RABV: Rabies lyssavirus
RPV: Rinderpest virus
RVFV: Rift Valley fever virus
SARS-CoV-2: Severe acute respiratory syndrome coronavirus 2
SBV: Schmallenberg virus
SHV: Suid alphaherpesvirus 1
SPoxV/GPoxV: Sheeppox/Goatpox virus
SVDV: Swine vesicular disease virus
USUV: Usutu virus
VSBV-1: Variegated squirrel Bornavirus-1
VSIV: Vesicular stomatitis Indiana virus
WNV: West Nile virus

## Ethics approval and consent to participate

For samples L02912 and L02913, animal experiments were evaluated and approved by the ethics committee of the State Office of Agriculture, Food safety, and Fishery in Mecklenburg – Western Pomerania (LALLF M-V: LVL MV/TSD/7221.3-1-067/18). For samples L02456 and L03827-30 as well as L03862-64 ethics approvals are documented in the publications as cited in Table 2, references 30 and 31, respectively.

## Consent for publication

Not applicable.

## Availability of data and materials

The bait set generated and tested in this study including the GenBank accession number of the respective virus genomes are provided in Additional file 1 (fasta format).

## Authors’ contributions

CW and MB conceived the study and wrote the manuscript, CW performed the experiments and analyzed the data. SC analyzed data, KS and UZ shared samples with known viral content. DH and AP helped with study design, discussed the results, and edited the manuscript. All authors read and approved the final manuscript.

## Acknowledgments

We acknowledge Brian Brunelle and Alison Devault (Arbor Biosciences) for professional help during bait design. We thank Andrea Aebischer Sandra Blome, Sandra Diederich, Martin Eiden, and Michael Eschbaumer for providing the proficiency test samples. We are grateful to Dennis Rubbenstroth and Arnt Ebinger for providing genome sequences; Patrick Zitzow for excellent technical support; Leonie Forth, Jan Forth, Conrad Freuling, Florian Pfaff, Pauline Santos, Jacob Schön and Adi Steinrigl for sharing sequencing libraries. Finally, we thank Arbor Biosciences for providing the predesigned SARS-CoV-2 bait set.

